# Microbial communities in *Ornithodoros phacochoerus* ticks: spatial structure and influence of African swine fever virus infectious status

**DOI:** 10.64898/2026.02.26.708205

**Authors:** Florian Taraveau, David Bru, Hélène Jourdan-Pineau, Elsa Rudo Pires Lameira, Carlos João Quembo, Mélanie Jeanneau, Maxime Duhayon, Antonia Andrade, Alberto Francisco, Jacinto Chapala, Thomas Pollet

## Abstract

*Ornithodoros phacochoerus* are nidicolous soft ticks of the *Ornithodoros moubata* complex of species known to be vectors of the African swine fever (ASF) virus. These *Ornithodoros* ticks depend on endosymbionts to produce essential nutrients necessary for their development. However, endosymbionts are only a part of the complex microbiota hosted by the tick. This microbiota often includes primary or secondary endosymbionts, commensal species from the environment, and, most of the time, some pathogens.

The present study was performed to understand the organization and spatial distribution of the microbiota of *O. phacochoerus*. One of the objectives was to investigate if the pathogen of interest (ASF virus) is involved in the organization of the microbiota through pathogen-induced dysbiosis or other interactions. For this purpose, 704 *O. phacochoerus* ticks were collected from two conservation areas in Mozambique. Sequencing was performed targeting the V3-V4 region of the 16S rRNA gene, and the resulting dataset was processed using FROGS to characterize the bacterial microbiota hosted by the ticks.

The results indicate that the microbiota of *Ornithodoros phacochoerus* contains very low bacterial diversity, with one primary endosymbiont (*Francisella*-like endosymbiont), one potential secondary endosymbiont (*Rickettsiella*), and very few environmental or pathogenic bacterial species. We found a clear spatial structure of the microbiota, with ticks from the same sampling site showing similar patterns. On the contrary, no association with the infectious status for African swine fever virus was detected, suggesting that this pathogen does not shape *Ornithodoros* microbial communities. Our results on tick – microbiota – pathogen – environment interactions in nidicolous soft ticks, showed patterns that differ from most hard tick studies.

## Introduction

Over the past two decades, next-generation sequencing methods led to a strong increase in the number of microbiota studies (Greay et al. 2018). The composition, distribution and then the role of the microbiota could thus be investigated in numerous organisms, including many arthropod species. In particular, blood feeding arthropods have received much attention due to their role as vectors of pathogens. This is the case for ticks for which many studies highlighted the complexity of tick microbial communities which include endosymbionts, commensals and pathogens. Endosymbiontic intracellular bacteria are maternally transmitted and provide their host with B vitamins which are not naturally brought by the strictly hematophagous nutrition (Duron *et al*., 2018; Bonnet *et al*. 2017). Those endosymbionts are also known to have an influence on fitness, reproduction (Guizzo *et al*. 2017; Li, Zhang, & Zhu 2018; Zhang *et al*. 2017; Zhong, Jasinskas, & Barbour 2007; Kurlovs *et al*. 2014), immunity, and vector competence (Narasimhan *et al*. 2014; Bonnet & Pollet 2021; Gall *et al*. 2016). Commensal bacteria are acquired from the environment and may differ from one individual to another. They are not transmitted vertically and their biological effects on ticks remain largely unknown, (Narasimhan & Fikrig 2015).

Interactions between pathogens and the tick’s microbiota turned out to be very diverse (Bonnet and Pollet 2021), ranging from synergies facilitating the infection (Narasimhan et al. 2014; Gall et al. 2016) to competition or protection against pathogens (Gall et al. 2016) or even pathogens reshaping the composition of the microbiota (Adegoke et al. 2020; Abraham et al. 2017). These interactions are particularly important to understand as ticks host complex microbial communities which will systematically be encountered by the pathogens. Moreover, it is now well established on hard ticks that microbial communities are very diverse, dynamic in both space and time (i.e. Carpi et al. 2011; Lalzar et al. 2012; Van Treuren et al. 2015; Lejal et al. 2021; Krawczyk et al. 2022). Spatial structuration in microbiome has mainly been demonstrated in tick pathogens (e.g. Joly-Kukla et al. 2024) and is driven by ecological factors such as the spatial distribution and density of reservoir hosts (Ostfeld et al. 2006).

Most of the studies available in the literature focus on hard ticks. Unlike most hard ticks, soft ticks are mostly nidicolous species found in environments with controlled temperature and humidity such as nests, burrows or caves. Their life cycle is characterized by a short blood meal, long life span, and several nymph stages (Vial 2009). *Ornithodoros phacochoerus* is a soft tick of the *Ornithodoros moubata* species complex. These *Ornithodoros* are vectors of the African swine fever (ASF) virus which is of key importance in veterinary medicine (Sánchez-Vizcaíno et al. 2015). African Swine Fever Virus (ASFV) causes haemorrhagic fever in susceptible infected swine, with mortality rates of up to 100% in naïve domestic pigs. *Ornithodoros* of the *O. moubata* complex are widespread in southern Africa. In the wild, they are commonly found in warthog burrows, feeding on warthogs and other animals that occupy the burrows (Vial 2009). Few studies are available in the literature for soft ticks microbiota, all focused on *Ornithodoros* species. Microbiota composition and diversity varied with organs (salivary glands and midgut) in *O. erraticus* and *O. moubata* (Piloto-Sardiñas et al. 2023), with tick stage and sex in *O. maritimus* (Gomard et al. 2021) and with feeding status in *O. brasiliensis* (Dall’Agnol et al. 2021). O. turicata presented a high number of bacterial taxa potentially due to the variety of hosts (Barraza-Guerrero et al. 2020). All those studies confirm the predominance of endosymbiotic bacteria. None of these studies investigated the spatial dynamics of soft tick microbiota.

In this study, we thus investigated the composition and organization of the microbiota of *Ornithodoros phacochoerus*. Ticks of all developmental stages were collected in two conservation areas in Mozambique. Previous studies allowed to investigate the genetic structure of these samples using microsatellite markers (Taraveau et al. 2024, 2025). In the work presented here, we analyzed the bacterial microbiota of these ticks to understand how it is organized based on geographical and biological parameters, and in relation to the ASF virus.

## Methods

### TiCK sampling

*Ornithodoros phacochoerus* ticks were collected in Mozambique from two different conservation areas: the Coutada 9 Game Reserve from the district of Macossa, Manica province, and the Gorongosa National Park from the district of Gorongosa, Sofala province. Ticks from Coutada 9 Game Reserve were collected from 83 different sites out of which 19 sites were selected for NGS analysis. Ticks from Gorongosa National Park were collected from three different sites which were all kept for NGS analysis. More details on the sampling sites and sampling methods are detailed in Taraveau et al. (Taraveau et al. 2024), map of sampling sites is available in **Supplementary figure S1**.

Two geographical relevant levels were studied: 1/ the parks (Coutada 9 Game Reserve and Gorongosa National Park), and 2/ the sampling sites within a park. The type of sampling site was also taken into consideration with two different types of sites: “Boulder” which were shaded areas beneath granite boulders, where wild herbivores, including warthogs, spent time resting, and “Burrows” which were holes in the ground that were primarily dug by aardvarks (*Orycteropus afer*) and used by warthogs to sleep and rear their young, thus escaping predation.

Tick DNA was shared with the Mozambique Institute of Agricultural Research (IIAM) under Material Transfer Agreement and in compliance with the Nagoya Protocol.

### DNA extractions

Between 26 and 51 ticks were extracted in each site for a total of 704 ticks. Pictures of the ticks were taken to measure tick size using ImageJ software (Abramoff et al. 2004). As described in the literature, ticks were washed in a 1% bleach bath for 30 seconds, then rinsed for 1 minute in three consecutive baths of Milli-Q water to eliminate cuticular bacteria for other downstream analyses (Binetruy et al. 2019). Ticks were then cut and crushed individually. DNA was extracted from the crushed tick homogenate, using the standard protocol from the DNeasy® Blood & Tissue genomic DNA extraction kit (Qiagen, Hilden, Germany). DNA extracts were finally eluted in 200 µl of elution buffer and stored at -20°C until further use (Taraveau et al. 2024). Negative controls were added at each step: washing, crushing, extraction, and amplification (see below) (Lejal et al. 2020).

### DNA amplification

DNA amplification was performed on the V3-V4 region of the 16S rRNA. Primer and index sequences were based on the literature for multiplexing numerous samples to study microbial communities (Kozich et al. 2013). Primers were forward 5’-GTGCCAGCMGCCGCGGTAA-3’, and reverse 5’-GGACTACHVGGGTWTCTAATCC-3’. For each primer, one of 22 forward index sequences (8 bp) and 33 reverse index sequences (8 bp) were used to multiplex 704 tick samples and 22 negative controls. All amplification, normalization and multiplexing steps were performed under sterile conditions using UV decontamination for 20 minutes and DNA removal solution on material and surfaces. The amplification mix consisted in 5 μL of DNA template, 1x NGS Galan Phusion mix (Thermo Fisher Scientific, France), 0.35 μM of forward primer, 0.35 μM of reverse primer, in a final volume of 50 μL. The PCR program was set as follow: 98°C for 3 min, then 35 cycles of 98°C for 15 sec, 55°C for 30 sec and 72°C for 30 sec, followed by a final extension step at 72°C for 10 min. Amplification negative controls were added in each amplification plates by replacing DNA template with 5 μL of nuclease-free water. Amplicons were migrated on Sybr Green 1% agarose gels to check amplification success.

The same samples were also amplified to investigate for the African swine fever infectious status of ticks. Detection was performed with quantitative PCR using the taqMan method on Roche Lightcycler^®^ 96 instruments (Roche Applied Sciences, Beijing, China). ASFV was detected using primers and probes targeting *ASFV VP72* as previously described (Tignon et al 2011). The actin gene from *Ornithodoros moubata* was amplified with *ASFV VP72* as a control for the assessment of DNA extraction success and to standardize the results with the tick size. Primer sequences were Forward: 5’-CCGGTATTGCCGACCGTATGC-3’, Reverse: 5’-CTCCCTGTCCACCTTCCAGC-3’ and Probe: 5’-CGAGAGGAAGTACTCCGTCTGG-3’ (Pereira De Oliveira, Vial & Le Potier 2022; Duron *et al*. 2018). A plasmid encoding both the VP72 and tick beta-actin genes was used at different dilutions to obtain a standard curve and determine the number of gene copies in each sample (Pereira et al. 2020). From this study, 14 ticks out of 704 were detected positive for the ASF virus.

### Preparation of the library and sequencing

SequalPrep™ Normalization Plate kit (Invitrogen, United States) was used to normalize amplicons at equimolar concentrations. The obtained concentrations were checked using Qubit dsDNA High sensitivity Assay kit (Invitrogen, United States) and amplicons were then pooled at equimolar concentration.

The equimolar mix was finally sequenced by the Genotoul GeT platform (Toulouse, France) using the MiSeq Illumina 2*300 bp chemistry.

### Data treatment and analysis

Sequencing results were analyzed using FROGS v4.1.0 (Escudié et al. 2018) on the Galaxy instance of the Genotoul bioinformatics platform https://vm-galaxy-prod.toulouse.inra.fr (The Galaxy Community 2022). A total of 12,560,070 unmerged reads were obtained from sequencing. Demultiplexing and primer trimming were performed by the sequencing platform. Pre-process was performed to merge overlapping pairs of reads with a 0.1 mismatch rate using **vsearch** v2.17.0 (Rognes et al. 2016) leading to keep 3,302,482 paired-end assembled reads. Amplicon sequencing variants (ASVs) were obtained using the swarm clustering method with an aggregation distance of 1 using **Swarm** v2.1.12 (Mahé et al. 2015). Chimeric sequences were removed using **vsearch**. ASV filters were applied based on cluster prevalence (cluster needed to be present in at least four samples), and cluster abundancy (abundancy of at least 5×10^−4^ % of the total of reads) (Bokulich et al. 2013). Thirty-four clusters were kept after filtering, affiliation of each ASV seed was then performed using RDPClassifier (Wang et al. 2007) and NCBI Blastn+ (Camacho et al. 2009) against the 16S SILVA Pintail100 138.1 reference database (Quast et al. 2013).

The 22 negative controls included in the study were used to evaluate which ASV were contaminants based on the method available in the literature for tick microbial datasets (Lejal et al. 2020). ASVs corresponding to more than 1% of the sequences obtained in a control group were immediately identified as contaminants (8 ASVs). ASVs for which the mean number of sequences in negative controls was above the contamination threshold of 0.02% of the total number of sequences for this ASV were considered as contaminants (7 more ASVs) (Galan et al. 2016; Lejal et al. 2020). All 15 ASVs identified as contaminant (**Supplementary table S2**) were investigated using NCBI Blast and the literature to evaluate the relevance of this identification (Salter et al. 2014). After validation, contaminants were filtered from the dataset, leading to keep 19 ASVs representing 2,871,371 sequences in total. Please note that the most abundant ASVs detected in our dataset (*Francisella*, and *Rickettsiella* which are part of *Ornithodoros* microbiota) did not show signs of cross contaminations over the 0.02% threshold toward the negative controls.

After filtration, 95 tick samples contained less than 500 sequences and were consequently eliminated (Lejal et al. 2021). When needed, normalization of the number of sequences per sample was performed by random sampling (rarefaction step) to keep 1000 sequences per sample. Quality of sequencing depth was evaluated based on rarefaction curves (**Supplementary figure S3**). For most samples, the ASV richness reached a plateau before 2000 reads suggesting that a deeper sequencing would not add more ASVs and that sequencing coverage was sufficient.

### Statistical analysis

All diversity indexes were evaluated on FROGS with the package **phyloseq** v1.38.0 (McMurdie and Holmes 2013). Four α-diversity indexes were evaluated: observed richness which considers the number of observed ASVs; Chao1 which considers the observed richness and an estimate of the number of unobserved species; Shannon which considers species abundance distribution; and Inverse-Simpson which evaluates the probability that two sequences taken at random in the sample came from the same species. Due to a significant effect of sequencing depth on three out of four indexes (ANOVA, Observed richness F-value = 205, p-value < 2×10^−16^; Chao1 F-value = 188, p-value < 2×10^−16^; Shannon F-value = 14, p-value = 2.2×10^−4^), α-diversity was evaluated after the rarefaction step. Results from α-diversity analysis were compared on R version 4.2.3 (2023-03-15) (R Core Team 2023) to test for the effect of different conditions: sampling site, sampling park, type of site, African swine fever virus infectious status, and tick size. Generalized linear models were used with the function **glm** with a gamma distribution. The best model was selected using the function **dredge** (R package **MuMIn** (Burnham and Anderson 2004)). Statistical comparisons were then performed using chi-squared test, and post-hoc analysis was performed using the function **emmeans** (Tukey HSD test). Sequencing depth was detected to influence alpha diversity measures, this was corrected using a rarefaction step to normalize to 1000 sequences per sample.

Four β-diversity indexes were evaluated to compare samples pair by pair: Jaccard index evaluates the fraction of species (qualitative) which is specific to each of the samples compared; Bray-Curtis index evaluates the fraction of the community (quantitative) which is specific to each of the samples compared; Unifrac index uses a phylogenetic tree to compare the fraction (qualitative) of the tree specific to each of the samples; Weighted-Unifrac index is the same as Unifrac but comparing the abundance (quantitative) in the tree that is specific to each of the samples. Significance of composition differences depending on different groups (sampling sites, ASF status, tick size) was evaluated using multivariate ANOVA analysis with the distance matrix. Key results were visualized using multidimensional scaling (MDS or PCoA) and heatmaps. For ASF status (positive versus negative) and tick size (large versus small), differential abundance analysis was performed to evaluate if there were variations in ASV abundance between two conditions. This was performed using **DESeq2** v1.34.0 with **DESeq2** normalization method (Love, Huber, and Anders 2014) with population as a confounding factor. As there were only 14 positive ticks for ASF virus, the influence of this parameter was specifically evaluated in site number 49 which contained 5 positive ticks out of 28 ticks. Clustering of the ticks from site 49 was performed using ward clustering and a tree based graphical representation to evaluate the similarity of the five ASF positive ticks.

## Results

### *Ornithodoros phacochoerus* microbiota composition

From the microbiota of the 704 analyzed ticks, 19 ASV were identified belonging to 6 different genera: *Francisella, Diplorickettsia, Anaplasma, Staphylococcus, Bacillus*, and *Brevibacterium* (**Figure 1**). It should be noted that although *Diplorickettsia* have been described as a new bacterial genus (Mediannikov et al. 2010), their taxonomic classification remains unclear. *Diplorickettsia* are taxonomically nested within the genus *Rickettsiella* (Leclercque & Kleespies 2012; Duron et al. 2016), we thus concluded in this study the identification *Diplorickettsia* corresponded to the genus *Rickettsiella*. Alone, the two most abundant genera represented 84.0% (*Francisella*), and 15.8% (*Rickettsiella*) of the sequences. Other genera represented respectively 0.073% (*Anaplasma*), 0.072% (*Staphylococcus*), 0.023% (*Bacillus*), and 0.020% (*Brevibacterium*) of the sequences. *Francisella* were detected in 100% of the samples, while *Rickettsiella* were detected in 63.9% of the samples. The other genera were detected in respectively 19.7% (*Staphylococcus*), 18.9% (*Anaplasma*), 12.6% (*Bacillus*), and 5.4% (*Brevibacterium*) of the samples.

**Figure 1.**
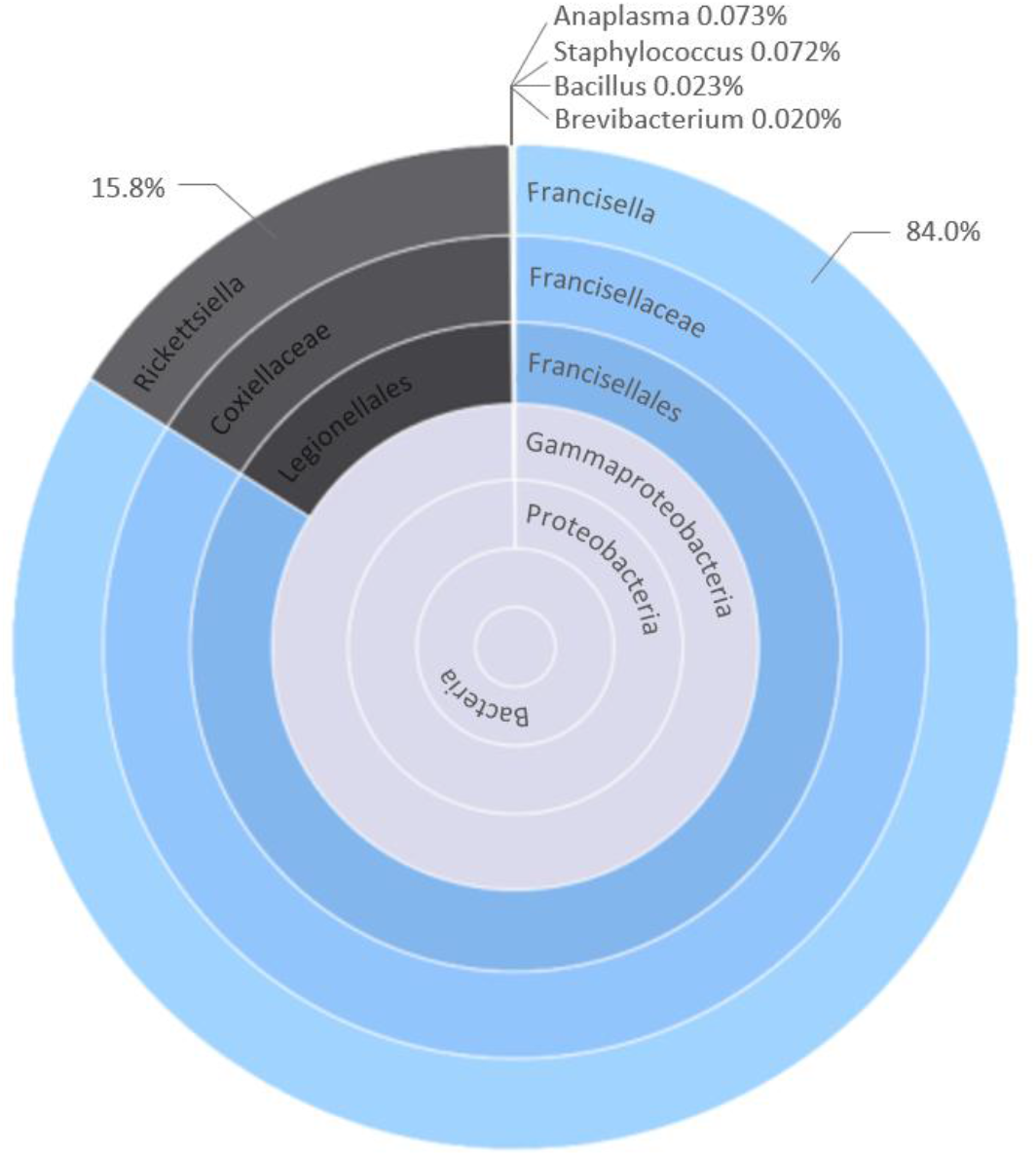
Pie chart of the genera detected in the microbiota of *Ornithodoros phacochoerus*. Percentage of total number of sequences are indicated.

Among the 19 ASV identified, several ones corresponded to the same genera and most of the time, the same species. Genera *Anaplasma* and *Brevibacterium* were represented by only one ASV each (2095 and 565 reads respectively). *Bacillus* and *Staphylococcus* were represented by two ASV each with similar numbers of sequences in both ASV (291 and 355 reads for *bacillus*; 875 and 1173 for *Staphylococcus*). *Rickettsiella* consisted of four ASV, with one ASV containing 98.5% of *Rickettsiella* sequences (445,857 reads for the main ASV, between 477 and 4702 reads for the others). *Francisella* consisted of nine different ASV, with one ASV containing 99.4% of *Francisella* sequences (2,382,126 reads for the main ASV, between 169 and 8216 reads for the others). A phylogenetic tree of ASV affiliations is presented in **Supplementary figure S4**. Mean observed richness was evaluated at 4.04 ASV per sample (SD=3.00, 95% CI [3.80; 4.29]), Chao1 at 4.62 ASV (SD=3.93, 95% CI [4.30; 4.94]), while mean Shannon index was evaluated at 0.37 (SD=0.32, 95% CI [0.34; 0.39]).

### Effect of geographical structure

Alpha diversity indexes were evaluated using **glm** models to take into consideration the effect of the geographical distribution (depending on the parc, the type of site, and the sampling site), ASF status (positive or negative) and tick size (small *ie* stage 1 nymph, medium *ie* stage 2 or 3 nymphs, or large *ie* stage 4 or 5 nymphs or adults).

In all analyses of α-diversity, the sampling site was the only parameter kept in the best model that fit the data (X^2^= 522, p-values < 2.2×10^−16^ for observed diversity, X^2^= 187, p-values < 2.2×10^−16^ for chao1, X^2^= 22, p-values < 2.2×10^−16^ for Inverse Simpson indexes, and X^2^= 117, p-values = 1.2×10^−15^ for Shannon index). All four indexes gave similar results concerning which populations presented less diversity: the five populations with the less diversity were sites number 58, 70, 71, 72, and 69. The sites with highest diversity values were sites number 3, 15, 49, G2, and G3 for Chao1 and observed diversity, and 15, 10, 16, 3, and 54 for Shannon and Inverse Simpson (p-value < 0.0001, Bonferroni correction for 210 tests). The type of site (burrows versus rocks) and the parc (Gorongosa versus Coutada 9) did not bring any amelioration to the model containing only the sampling site. However, except for site number 5 which contained only one individual after normalization, all other rock sites (sites number 3, 5, 10, 15, 16, and 49) were among the top ten sites with highest diversity values out of 22 sites (**Figure 2** and **Figure 3**).

**Figure 2.**
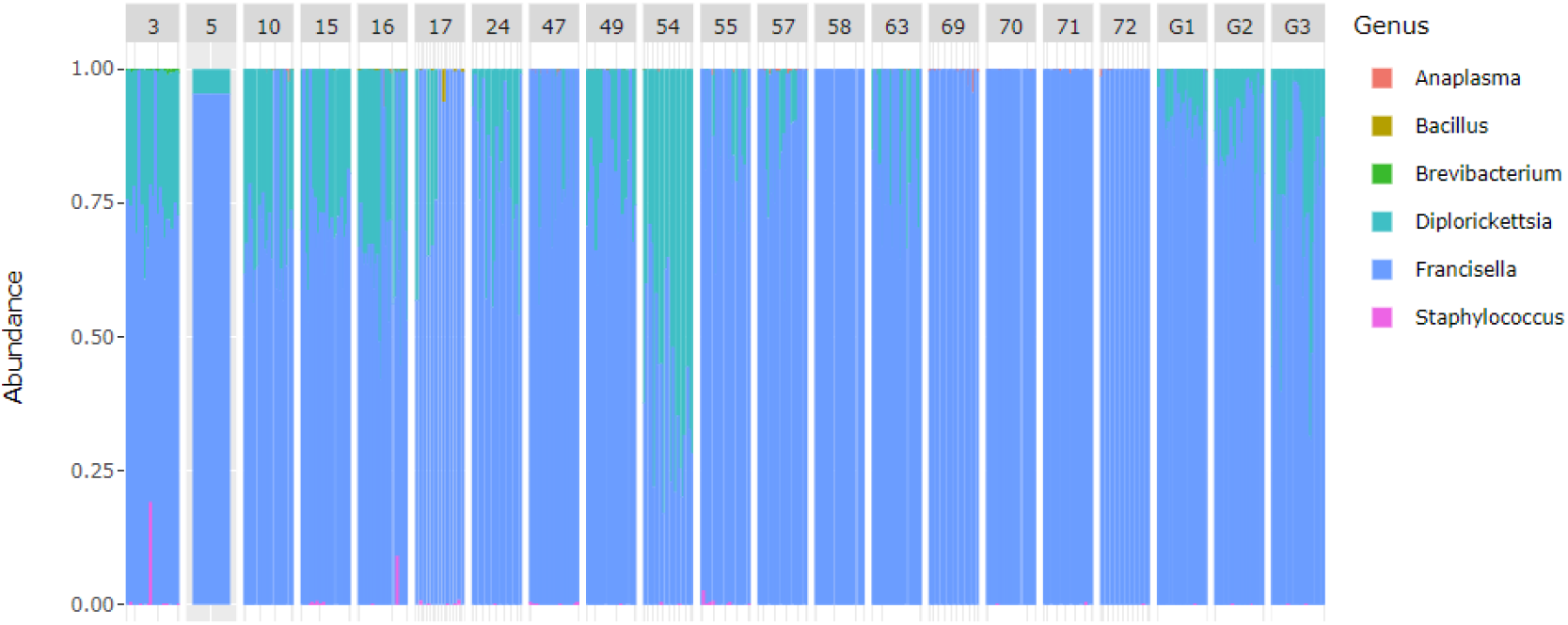
Observed diversity in each sampling site after rarefaction step.

**Figure 3.**
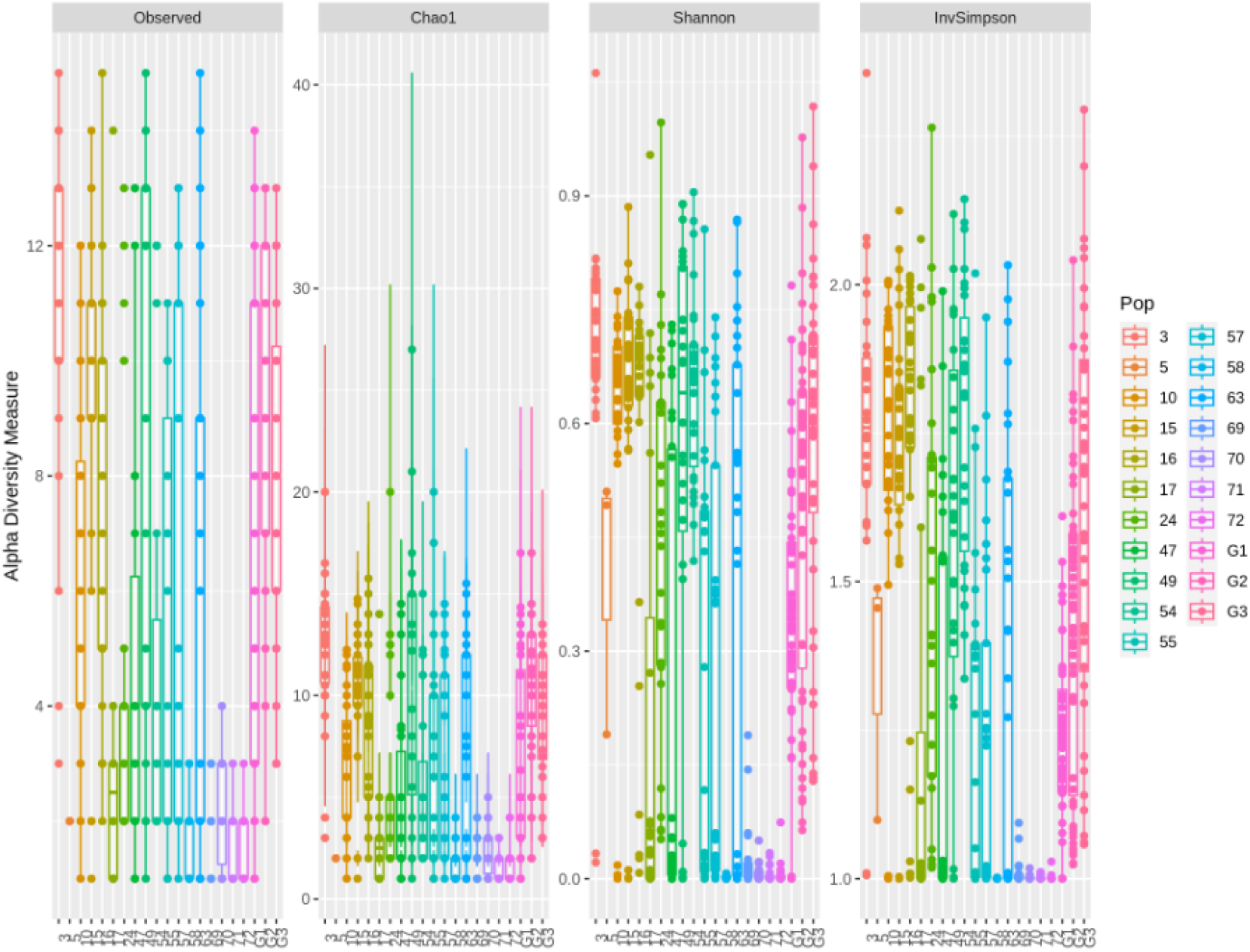
Four different α-diversity indexes depending on the sampling site (Pop).

Using pairwise comparisons between pairs of ticks with β-diversity indexes, the sampling site explained respectively 34% (Jaccard, p-value < 1×10^−4^), 48% (Unifrac, p-value < 1×10^−4^), 56% (Weighted-Unifrac, p-value < 1×10^−4^), and 57% (Bray-Curtis, p-value < 1×10^−4^) of the variations in diversity, making it the most significant parameter influencing β-diversity. Bray-Curtis visualization of ASV distribution is presented in **Figure 4** and highlight several profiles of microbial composition, including several sites without any *Rickettsiella* (sites 69, 70, 71, 72) and one site containing most *Brevibacterium* and *Staphylococcus* sequences (site 3). Moreover, several secondary ASVs for *Francisella*-like endosymbionts also appeared to depend on the sampling site in a pattern similar to the distribution of Rickettsiella (**Figure 4A**), most of them were not detected in the sites 69, 70, 71 and 72.

**Figure 4.**
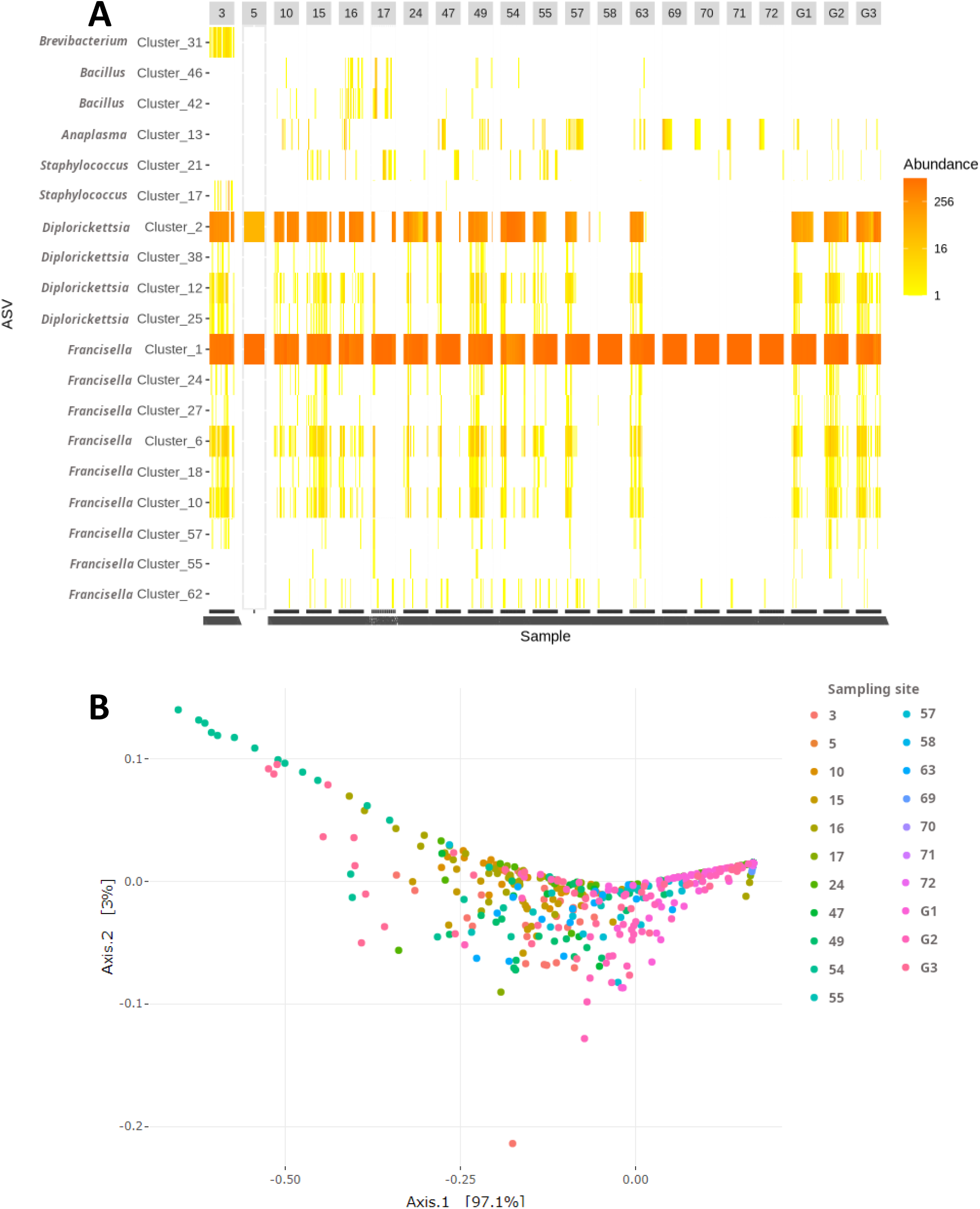
Visual representation of Bray-Curtis β-diversity index according to the sampling site. **A.** Heatmap of clusters’ distribution between samples. **B**. PCoA of ticks from the different sampling sites.

### Link with infectious status for African swine fever virus

After normalization, 11 out of 14 positive ticks for ASF virus remained in the dataset. These ticks came from five different sites with five ticks from site 49. With the strong effect of the sampling site on tick microbial diversity, it was decided to analyze separately the situation from site number 49 (28 ticks) to evaluate links between ASF virus infection and microbiota composition. As previously observed over the full dataset, no difference was detected for α-diversity indexes between ASF positive and negative ticks (ANOVA; Observed, p-value = 0.77; Chao1, p-value = 0.65, Shannon, p-value = 0.82, Inverse-Simpson, p-value = 0.59). Similarly, no difference was detected for β-diversity indexes (Multivariate ANOVA; 9999 permutations; Bray-Curtis, p-value = 0.59; Jaccard, p-value = 0.94, Unifrac, p-value = 0.83, Inverse-Simpson, p-value = 0.68). Ward clustering was applied to samples from site 49 to evaluate which ticks are similar in term of microbial composition, results are presented in **Figure 5**.

**Figure 5.**
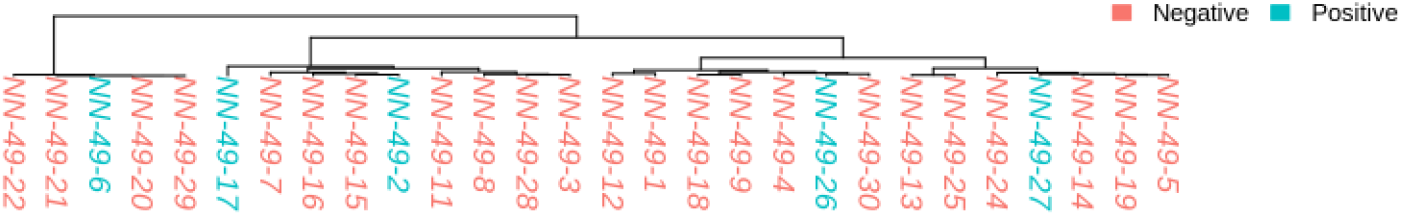
Clustering tree after Ward clustering of ticks from sampling site number 49. ASF positive ticks are indicated in blue, ASF negative ticks are indicated in red

Differential abundance analysis was performed to check if any ASV would be present in different abundances between ASF positive and negative ticks, with the sampling site as a confounding factor using negative binomial generalized linear model: no such ASV was detected.

### Effect of life stage

The number of reads obtained after sequencing for each tick was strongly correlated with the tick size (Spearman correlation test, rho = 0.30, p-value = 4.15×10^−15^). Especially stage 1 nymphs (small ticks) presented significantly less reads than medium or large size ticks (Kruskal test, chi^2^= 167.96, p-value < 2.2×10^−15^; Dunn test “Large – Small” corrected p-value = 1.72×10^−29^, “Medium – Small” corrected p-value = 6.19×10^−34^).

When comparing large ticks versus small ticks with DESeq2 (considering the sampling site as a confounding factor), it appeared that five clusters of *Francisella* and three clusters of *Rickettsiella* were less abundant in small nymphs than in large nymphs and adults (**Supplementary figure S5**). No such effect was detected for the other clusters, including clusters 1 and 2 which were the main clusters of *Francisella* and *Diplorickettsia* sequences respectively.

## Discussion

In this study, we investigated the microbiota of a nidicolous soft tick (*Ornithodoros phacochoerus*) which is found in warthog burrows. We described the microbial communities harbored by this tick and their organization between spatially separated sites, and we assessed whether this organization was altered by the presence of a pathogen, the African swine fever (ASF) virus.

Based on the analysis of the microbiota of the 704 ticks, only six bacterial genera were identified, with a mean Shannon index of 0.37. These results suggest that the microbiota *Ornithodoros phacochoerus* is very limited in term of species diversity. Compared to hard tick species, up to ten times more ASVs and bacterial genera could be found in most tick species (Heise, Elshahed, and Little 2010; Maldonado-Ruiz et al. 2021; Portillo et al. 2019; Lejal et al. 2021). It is important to keep in mind that comparing microbial diversity from one study to another is an arduous task. Here, we applied very stringent filters to the dataset studied. Ticks were washed with bleach to remove cuticular bacteria (Binetruy et al. 2019), and multiple negative controls were added to clean our dataset from as many contaminants as possible (Lejal et al. 2020). The low diversity observed did not appear to be related to insufficient sequencing depth, as a plateau was reached for the rarefaction curves of most samples and the Chao1 profile was very similar to observed richness suggesting that most ASV had been detected. Few studies are available in the literature for soft ticks. The diversity found here was very similar to that previously found in one laboratory strain of *Ornithodoros moubata* (Duron et al. 2018). However, other *Ornithodoros* species could present much higher diversity, such as observed in *O. turicata* (Barraza-Guerrero et al. 2020), *O. brasiliensis* (Dall’Agnol et al. 2021), *O. maritimus* (Gomard et al. 2021) or *O. erraticus* and another laboratory strain of *O. moubata* (Piloto-Sardiñas et al. 2023). Differences in protocols (no washing of cuticular bacteria, and/or no elimination of sequences from negative controls) make all these studies difficult to compare in terms of global diversity. Yet, even with protocol differences, the diversity observed in *O. phacochoerus* remained surprisingly low. It is possible that the restricted environment (warthog burrow) in which the ticks spent most of their life was not a great source of diverse environmental bacteria. Most bacterial communities are obtained from the environment and from maternal transmission (Narasimhan and Fikrig 2015). Being a nidicolous species, *O. phacochoerus* has a limited activity with almost no movements in the environment, except when searching for its host. This could lead the ticks to acquire almost no environmental bacteria, and to maintain mostly maternally inherited symbionts which are transmitted from one generation to another. Interestingly, in *Ornithodoros* soft ticks, the midgut possesses an antibacterial activity, amplified after blood meal, which could also participate to maintain low bacterial diversity in the midgut as detected here (Nakajima et al. 2003).

Going into the detail of the genera identified in *Ornithodoros phacochoerus*, two genera represented the vast majority of the obtained sequences: *Francisella* and *Rickettsiella*. The detection of *Francisella* in *O. phacochoerus* is not surprising as it has already been identified as a major primary endosymbiont of ticks of the *O. moubata* species complex (Duron et al. 2018), the species complex to which *O. phacochoerus* belongs. *Francisella*-like endosymbionts have been well identified to have a nutritional role, synthetizing B vitamins, an essential nutrient not brought by the strictly hematophagous alimentation of ticks. *Coxiella*-like endosymbionts and several other bacterial species have been identified as capable of assuring this nutritional role (Duron and Gottlieb 2020), including in some *Ornithodoros* species such as *Ornithodoros brasiliensis* (Dall’Agnol et al. 2021). As a consequence, alternatives to *Francisella* may have occurred, although they were not likely, based on previous observation from closely related species. It was expected that the primary endosymbiont associated with B vitamin synthesis should be present in all individuals. This was confirmed here with 100% of the ticks having *Francisella*. Interestingly, nine *Francisella* haplotypes were detected (with one largely predominant). This could be the result of sequencing errors leading to false ASVs that differ only slightly from one another and have very few reads. Such ASVs could be detected in the affiliation tree (**Supplementary figure S4**) where the clusters are very close from one another with only a few substitutions in the DNA sequence. Conversely, it may also indicate that several strains of the same symbiont coexist in ticks, probably as a result of genetic drift. These different strains could be complementary (production of different nutrients) or more adapted to certain conditions (differences in temperature, humidity, availability of the hosts…). Several *Francisella* haplotypes had already been detected in other tick species (Buysse et al. 2022). However, the fixation of a new haplotype occurred only following strict and long-term geographic isolation. In the case of our dataset, such a barrier only existed between the two conservation areas (Coutada 9 versus Gorongosa, Taraveau et al. 2025) but no differences in ASVs composition was detected between the two parks.

Interestingly, some laboratory strains of *O. moubata* also showed 100% infestation with a bacteria of the genus *Rickettsia* (Taraveau et al. 2023). Such *Rickettsia* were also found in other arthropods, and in various genus of both soft ticks and hard ticks (Duron et al. 2017). In *O. phacochoerus*, we observed a high prevalence of *Rickettsiella* (63.9% of the ticks), with abundance levels sometimes exceeding Francisella and highly varying across tick populations. This indicates that they are probably facultative for the tick’s life cycle, a feature of secondary symbionts. *Rickettsiella* are known to be frequently part of *Ixodes* and some *Ornithodoros* microbiota, with maternal transmission occurring, and a prevalence in up to 90% of the ticks (Hodosi, Kazimirova, and Soltys 2022; Garcia-Vozmediano et al. 2022; Duron et al. 2015). In addition, a recent study revealed that certain genomes of *Rickettsia* and *Rickettsiella* strains have lost the integrity of their virulence genes, but have acquired homologs of *Wolbachia cif* genes which are known to induce cytoplasmic incompatibility in various arthropods (Floriano et al. 2025). One of these *Rickettsiella* strains was isolated from four *O. phacochoerus* ticks sampled in Coutada 9 Game Reserve. This suggests that the *Rickettsiella* strains detected in our samples might use a similar molecular mechanism to manipulate the reproduction of their arthropod hosts. Further experiments would be needed to confirm this role.

In term of tick-borne pathogens, one well-documented pathogenic genus was detected in our *O. phacochoerus* ticks: *Anaplasma. Anaplasma* are intracellular bacteria mostly found in *Ixodes* ticks and other hard ticks (Moraga-Fernández et al. 2023). Eight *Anaplasma* species were reported so far, causing persistent infections (anaplasmosis) in ruminants, humans, domestic and wild animals. Previous observations reported the presence of *Anaplasma* in *Ornithodoros* soft ticks, in *O. lahorensis* in China with a prevalence of 1.5% (Li et al. 2023), and in *O. moubata* in Zambia in 102/124 pools of 3-5 ticks from warthog burrows (Qiu et al. 2021). No complete analyses of vector competence have been performed on *Ornithodoros* so far, however artificial feeding on *O. fonsecai* and *O. brasiliensis* seems to indicate that *Ornithodoros* ticks can acquire *Anaplasma* through the blood meal and maintain it through transstadial transmission (Castro-Santiago et al. 2023). With all this information, it is probable that the *Anaplasma* detected here came from vertebrate hosts on which the ticks took a blood meal. Further studies would be needed to identify more precisely the pathogenicity of the strain detected here and the risk of transmission by *Ornithodoros* soft ticks.

Afrotropical *Ornithodoros* are well known vectors of *Borrelia* bacteria responsible for relapsing fevers (Cutler 2015). Yet, no *Borrelia* were detected in our dataset. In the literature, rodents and domestic animals are identified as hosts for relapsing fever *Borrelia*, but more analyses are needed on the role of wildlife (Lopez, Hovius, and Bergström 2021). Such *Borrelia* were previously detected in Zambia in *O. moubata* and *O. porcinus* ticks from warthog burrows with 43/182 positive pools of 2-8 ticks (Qiu et al. 2023). As for many pathogens, it is possible that the number of ticks sampled here was not sufficient to detect the presence of the *Borrelia*. However, with no *Borrelia* detected out of 704 ticks, if this bacteria is present, it would be in very low prevalence compared to the results obtained in Zambia. From all these data, it is possible to say that the risk associated with *Borrelia* in Coutada 9 Game Reserve is probably very low.

Finally, the last three genera detected were *Brevibacterium, Staphylococcus*, and *Bacillus*. Based on the affiliation blast and previous literature, these genera probably represent environmental bacteria that ended up in the digestive tract of ticks. *Brevibacterium* have been previously detected in *O. maritimus* (6.6% of the sequences) and other arthropod species (Gomard et al. 2021). No specific role or pathogenicity is known for these bacteria, although a role in blood digestion has been hypothesized due to their presence in several blood feeding arthropods. *Staphylococcus* was also found in *O. maritimus* with stable positive interactions with *Brevibacterium* at all stages of development of the tick (Gomard et al. 2021) both in *O. brasiliensis* (Dall’Agnol et al. 2021) and in *O. tholozani* (Karim et al. 2017). These studies suggest that *Staphylococcus* bacteria probably came from the skin and fur of vertebrate hosts. For *Bacillus*, they were previously found in *Ixodes ricinus* where they were detected only in a few ticks (Lejal et al. 2021). In this study, *Bacillus* strongly participated in the correlation networks. They were positively correlated with *Rickettsiella* and *Anaplasma* and other tick-borne pathogens and environmental bacteria. In a study on *Rhipicephalus microplus, Bacillus* were associated with the presence of *Theileria* in ticks (Adegoke et al. 2020). Both studies suggest that *Bacillus* could be associated with tick-borne pathogens. This could be due to their ability to survive in unfavorable physiological conditions that could be the result from competition between environmental microbiota and pathogens. Overall, the presence of these environmental bacteria must not be overlooked as it could be linked with environmental differences or tick-borne pathogens.

In summary, the microbiota of *O. phacochoerus* showed low diversity, with an obligate primary endosymbiont probably involved in nutrional symbiosis and a facultative secondary endosymbiont whose role in cytoplasmic incompatibility needs to be confirmed. We also described a few environmental species, and one potentially pathogenic bacterium. Such results may be related to the environment in which these ticks are found. As nidicolous ticks, *O. phacochoerus* ticks spend most of their lives in warthog burrows, waiting for a potential host to feed on. The blood meal is short (less than two hours), the ticks can live up to 20 years, and can remain in anfractuosities for several years between each blood meal (Vial 2009) without moving as they even rely on environmental humidity for their hydration. Such limited environment and short contact with vertebrate hosts could prevent *Ornithodoros* ticks from being exposed to multiple environmental bacterial strains, limiting the chances of colonization at each generation by commensal bacteria, whereas endosymbionts are transmitted maternally, and *Anaplasma* would come from vertebrate hosts. Interestingly, ticks collected from resting sites rather than burrows appeared to have higher alpha diversity indexes. Resting sites under granite boulders are open muddy areas, used by several mammal species to rest. Such areas are less isolated and offer more feeding opportunities for ticks than underground burrows, potentially allowing the ticks to encounter more commensal bacteria (Narasimhan and Fikrig 2015).

In *Ornithodoros phacochoerus*, larvae molt into stage 1 nymphs without taking any blood meal in about twenty days (Taraveau et al. 2023). This means that stage 1 nymphs did not feed yet on any vertebrate hosts. The microbiota that they contain most probably come from maternal transmission or direct environmental source (and not from the host skin for example). *Francisella* and *Rickettsiella* are already well known to be maternally transmitted (Duron et al. 2017). The frequent presence of *Rickettsiella* in stage 1 nymphs confirms the hypothesis of a probable maternal transmission for this bacteria. Interestingly several haplotypes from both *Francisella* and *Rickettsiella* were found to be less abundant in nymph 1 ticks than in adults. Two hypotheses could be made. 1/ this is the result of a sequencing bias, nymph 1 presenting much less biological material, leading the sequencing depth to be lower in these samples and, consequently, to lack several ASVs. This is supported by the correlation between tick size and the number of reads obtained for the tick. 2/ this is a biological result and only one/a few *Francisella* and *Rickettsiella* ASVs are maternally transmitted. This would mean that the other ASVs detected are the result of mutations and are mostly not transmitted to the offspring. Endosymbionts are particularly exposed to mutations and genetic drift as the loss of non-essential genes can include losing DNA repair genes (Moran, McCutcheon, and Nakabachi 2008). As symbiont populations bloom after a blood meal (Taraveau et al. 2023), there are more divisions in symbiont populations from adult ticks than from nymphs. Consequently, there are more opportunities for mutations and the emergence of secondary ASVs.

One objective of this study was also to evaluate whether the microbiota of *Ornithodoros phacochoerus* was spatially structured. Based on the diversity indicators evaluated here, it appeared that spatial distribution was the main factor influencing microbial composition. The sampling site significantly influenced all diversity indexes tested. Differences between two separated conservation areas (Coutada 9 versus Gorongosa) were indistinguishable from the effect of the site in the models. It is interesting to note that *Ornithodoros phacochoerus* populations are genetically structured at the scale of the sampling site according to population genetics (Taraveau et al. 2025). Population structure was characterized by very limited exchanges of ticks at long distances but many exchanges at short distances. In this context, it is possible that when secondary members of the microbiota are not present in an area (as for *Rickettsiella* in sites number 58, 69, 70, 71, and 72), little to no external import would introduce it among the ticks. This could participate to the limited number of secondary symbionts maintained in *O. phacochoerus*. Similar geographical patterns were observed in other species. In *Ixodes ricinus*, maternally inherited symbionts were found to be unequally distributed geographically, with ticks from close areas presenting similar microbiota (Krawczyk et al. 2022; Van Treuren et al. 2015). This distribution was driven by geographic distances rather than randomly which is probably also what is happening in *O. phacochoerus*. Within a site, *Ornithodoros* shared quite similar microbiota. As one site does not seem to correspond to one family according to population genetics (Taraveau et al. 2025), it can be expected that the microbiota was shaped over several generations of ticks, leading to a homogenate pattern within a geographical sector. In this matter, it is also imperative to acknowledge the potential role of tick reproduction manipulation by some members of the microbiota, including *Rickettsiella* (Floriano et al. 2025), as cytoplasmic incompatibility can strongly influence distribution patterns. Further analyses are required to achieve a comprehensive understanding of this influence on the spatial distribution of tick microbiota.

The literature is full of studies investigating the link between microbiota and tick-borne pathogens (Bonnet and Pollet 2021). Effects described can vary from competition for resources (Gall et al. 2016), facilitation of the passage of the intestinal barrier (Narasimhan et al. 2014), to no effect at all. NGS studies do not allow to investigate if microbial species participate in the infection. However, correlations between microbial composition and infectious status could lead to make hypothesis on the role of some members of the microbiota or to suspect pathogen induced reshaping of the microbial communities (Adegoke et al. 2020). The main pathogen of interest for *O. phacochoerus* is the African swine fever (ASF) virus which is of great importance in veterinary medicine. In this study however, no significant correlations were found when comparing ASF virus positive and negative ticks, either on the entire dataset or when zooming on an infected site. This dataset only presented a few positive ticks and ideally more samples would be needed to increase the strength of such analyses. However, these results highlight some interesting hypothesis. First, as previously mentioned, *O. phacochoerus* presents a very limited microbiota, with almost no environmental species in the digestive tract. It is possible that ASF virus encounter no competition during tick infection due to this lack of diversity. Such phenomenon was described in the tick *Ixodes scapularis* where the infection with *Anaplasma phagocytophilum* initiated a dysbiosis of the microbiota by altering biofilm formation, dysbiosis which then facilitated the passage of the gut barrier by the pathogen (Abraham et al. 2017). This should need to be tested in *O. phacochoerus* by infecting ticks with a normal or modified microbiota to investigate differences in vector competence. Second, it seems probable that ASF virus is not strongly shaping the tick microbiota in the wild. We can hypothesize that stronger titers of ASF virus would disturb endosymbiont populations but this may then quickly lead to the death of the tick. Such tick death has been observed following virus infection with high titers (Rennie, Wilkinson, and Mellor 2000), but link with microbial communities needs to be investigated. Finally, in an endemic area for the virus, it is possible that frequent infections permanently shape the microbiota, leading to keep only bacteria which were not down-selected by the virus, and finally leading to the current microbial composition. However, such hypothesis remains very unlikely based on the low prevalence of ASF virus in Coutada 9 Game Reserve and the fact that only a few burrows are positive for the virus. Overall, more investigations are needed before drawing conclusion for the link between *O. phacochoerus* microbiota and African swine fever virus.

## Author contributions

FT collected the samples in the field, performed the experiments, analyzed the data, and wrote the manuscript. DB performed the experiments, and analyzed the data. HJ-P obtained the funding, coordinated the study, collected the samples in the field, and performed the experiments. EL performed the experiments, and analyzed the data. CQ obtained the funding, coordinated the study, provided the material, and collected the samples in the field. MJ, and MD collected the samples in the field, and performed the experiments. AA performed the experiments. AF and JC collected the samples in the field. TP obtained the funding, conceived and coordinated the study, performed the experiments and analyzed the data. All authors read and corrected the final manuscript.

## Acknowledgements

The authors would like to thank the students from the IIAM who helped for DNA extractions, Edna Manuel Meque, Isaque António Francisco, Matias José Dina, Neidy Alfredo Lopes, Albertina Jaime Chabissa and Luisa Duarte Madeira, as well as Estelle Pineau who helped for tick sampling and DNA extractions. We would like to thank Mokore and Western camps which provided help and shelter during sampling in Coutada 9 Game Reserve. We would like to thank Gorongosa National Park for permission to conduct research (scientific permit PNG/DSCI/C235/2022), we are grateful for the staff for their logistical support.

We are grateful to the Genotoul bioinformatics platform Toulouse Midi-Pyrenees and Sigenae group for providing help and storage resources thanks to their Galaxy instance (https://vm-galaxy-prod.toulouse.inra.fr), and to the Genotoul GeT-PlaGe plateform where the sequencing was performed. We would also like to thank Björn Rotter from GenXPro with whom we exchanged about sequencing methods.

Many thanks to FROGS support team for their help with the use of FROGS. We would especially like to thank Géraldine Pascal and Lucas Auer for the great formation and resources that they provided for FROGS users and the precious help and advice that they gave us.

Special thanks to Charlotte Joly Kukla for the discussions and scripts about NGS results analysis and for trying together to understand how to use FROGS.

## Data, scripts and codes availability

Datasets will be available online shortly after publication: https://doi.org/10.18167/DVN1/LGQFTI

## Supplementary material

**Supplementary figure S1:**
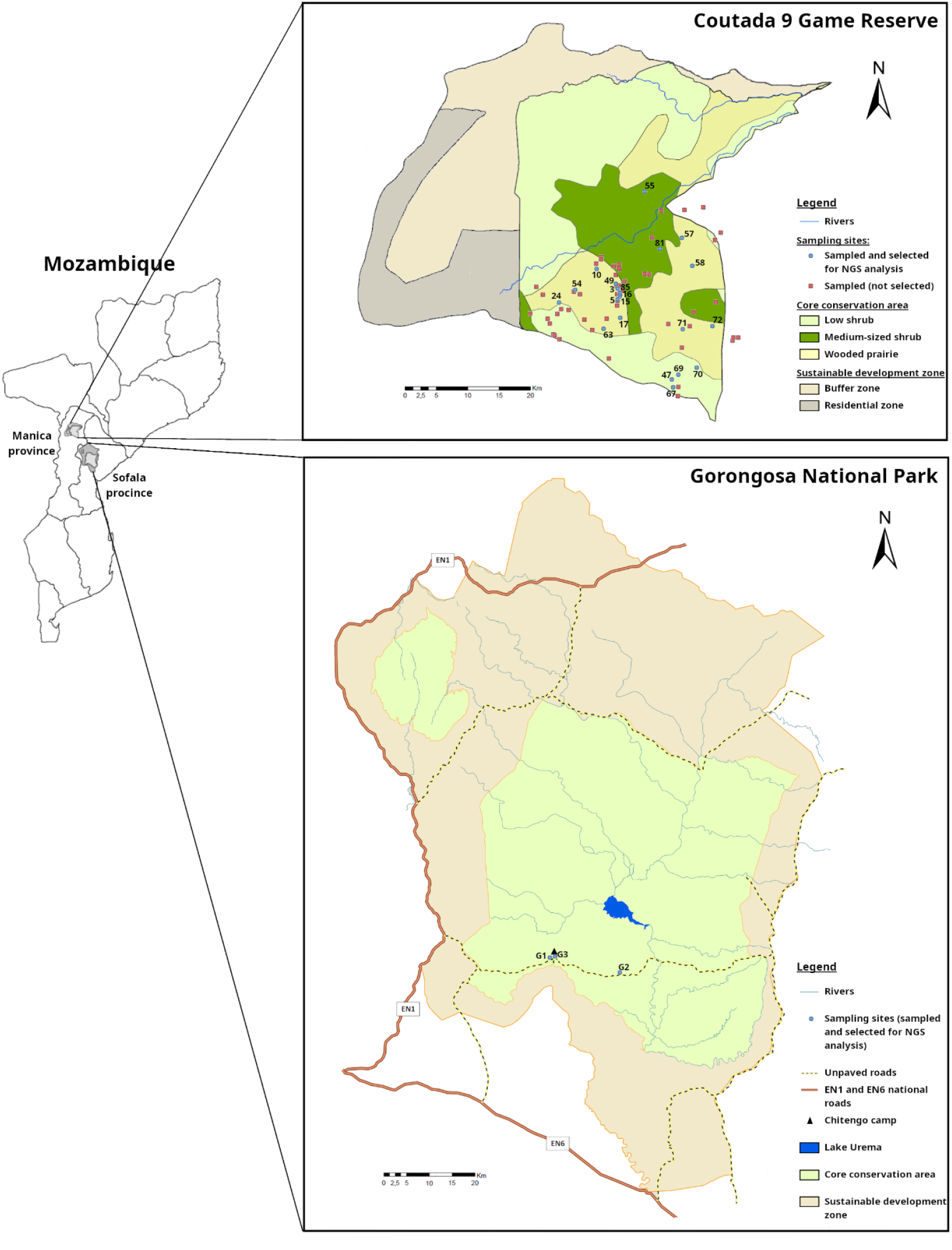
Location of sampling sites for *Ornithodoros phacochoerus* in Coutada 9 Game Reserve and Gorongosa National Park, Mozambique (Taraveau et al. 2025). For the sampled sites which were used in population genetics and/or NGS analysis (indicated with blue spots), the identification number of the site is indicated on the map.

**Supplementary table S2:**
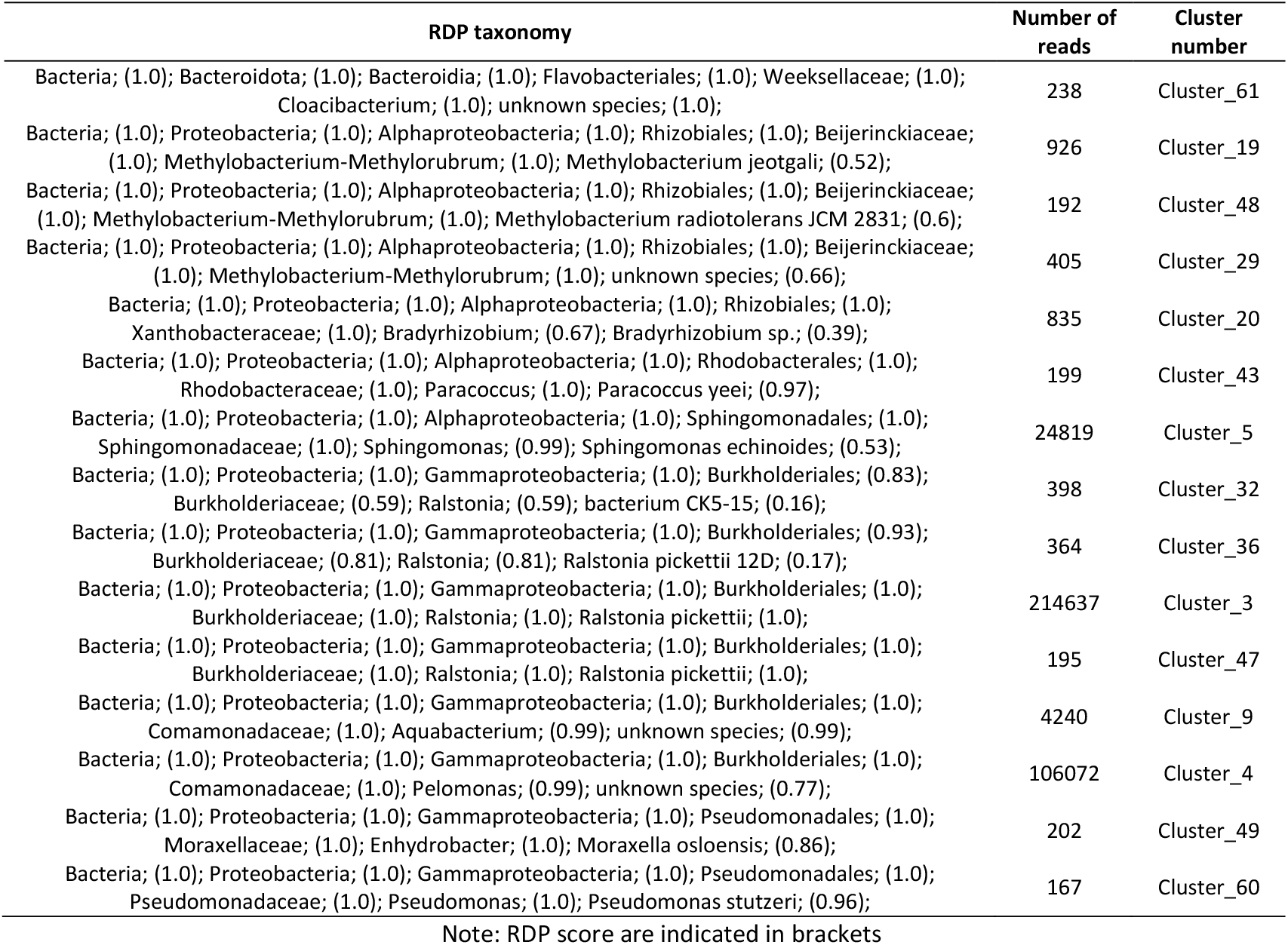
ASV identified as contaminants.

**Supplementary figure S3:**
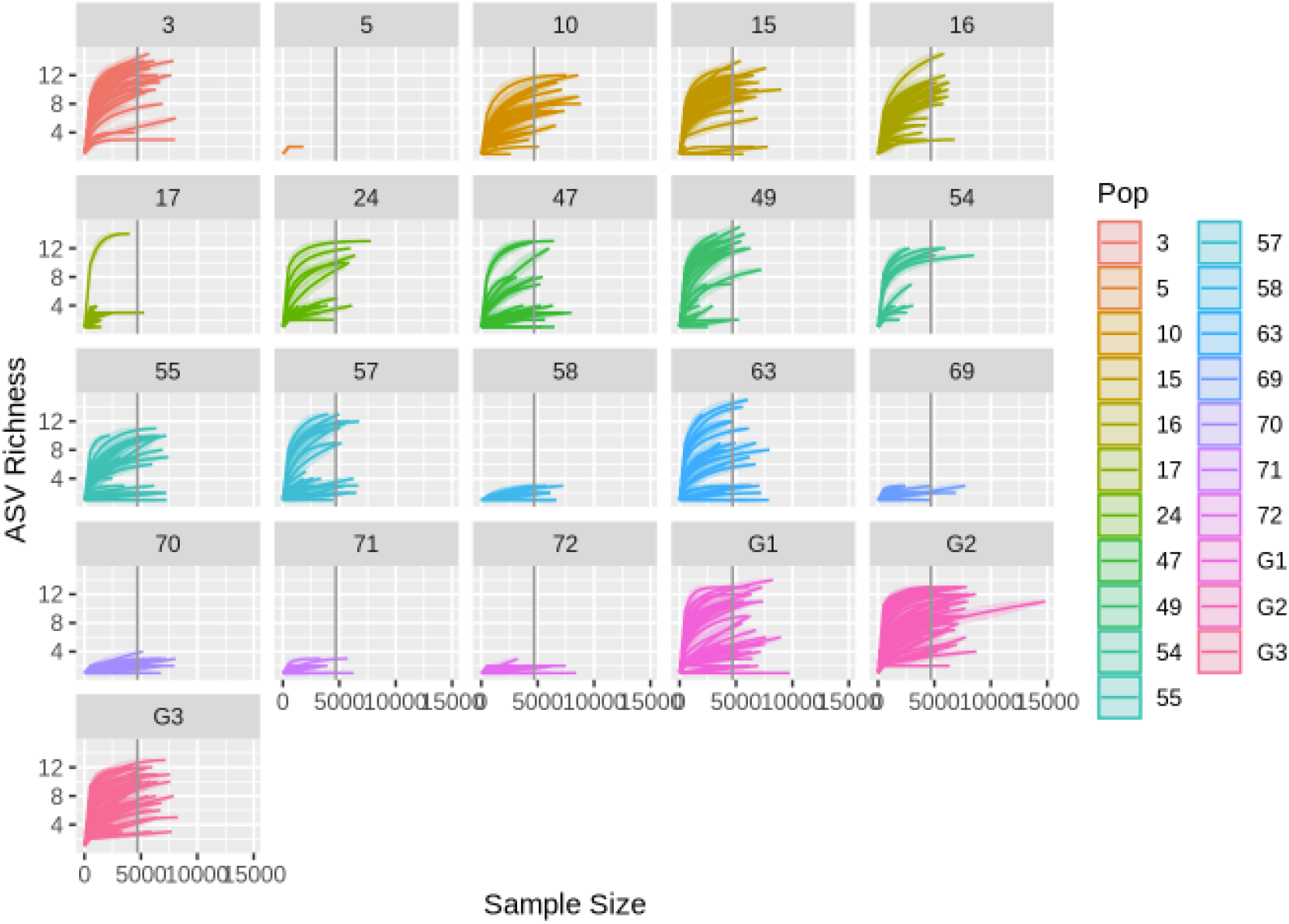
Rarefaction curves of samples colored by sampling sites (Pop)

**Supplementary figure S4:**
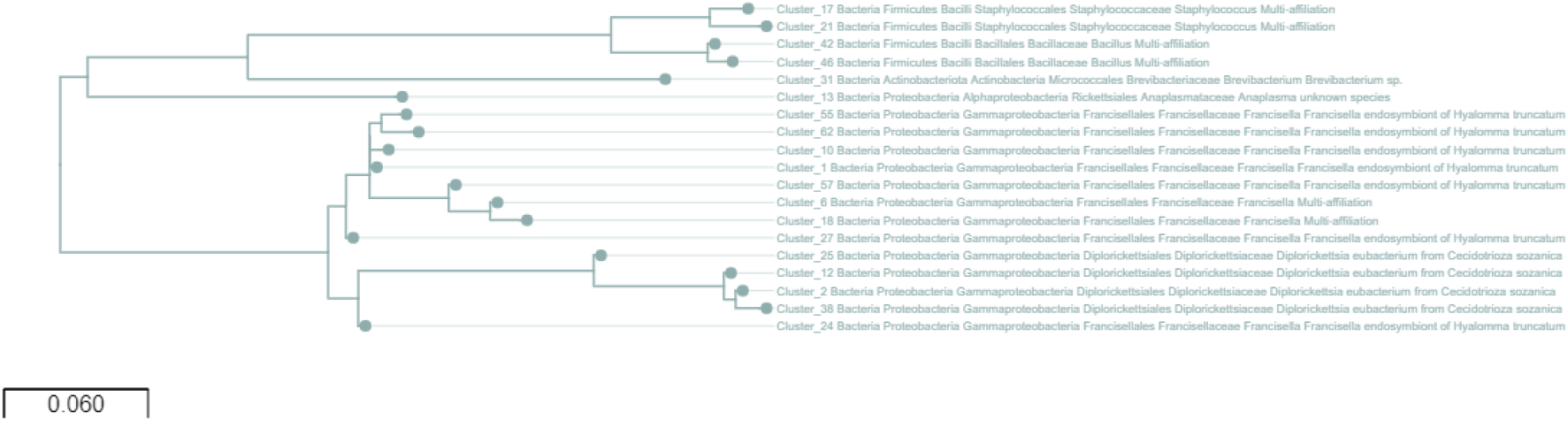
Phylogenetic tree based on the affiliations of the ASV detected. Multiple alignement performed using MAFFT, tree created with FastTree *via* FROGS.

**Supplementary table S5:** Differentially abundant ASV between small and large ticks.

| Cluster ID | Genus | Fold-change (log) | p-value |
| --- | --- | --- | --- |
| Cluster_6 | *Francisella* | -4.87 | $2.67 \times 10^{-16}$ |
| Cluster_10 | *Francisella* | -4.14 | $1.13 \times 10^{-14}$ |
| Cluster_12 | *Diplorickettsia* | -4.23 | $5.10 \times 10^{-14}$ |
| Cluster_18 | *Francisella* | -3.04 | $7.07 \times 10^{-10}$ |
| Cluster_25 | *Diplorickettsia* | -2.77 | $3.93 \times 10^{-7}$ |
| Cluster_24 | *Francisella* | -2.60 | $2.40 \times 10^{-6}$ |
| Cluster_27 | *Francisella* | -2.18 | 0.000224 |
| Cluster_38 | *Diplorickettsia* | -1.65 | 0.00475 |

Note: Negative fold-change implies that the ASV is more abundant in large ticks than in small ticks. P-values after Benjamini Hochberg correction.

## Conflict of interest disclosure

The authors of this preprint declare that they have no financial conflict of interest with the content of this article. Thomas Pollet is recommender for PCI Microbiology, Hélène Jourdan-Pineau is recommender for PCI infections and PCI EvolBiol.

## Funding

This study was supported and financed by: 1) the Ecology and Evolution of Infectious Diseases Program, grant no. 2019-67015-28981 from the USDA National Institute of Food and Agriculture, in a project entitled “unraveling the effect of contact networks & socio-economic factors in the emergence of infectious diseases at the wild-domestic interface” (https://www.asf-nifnaf.org/), 2) the Agence Nationale de la Recherche (ANR, France, ref. ANR-25-CE02-7068 CYTOTICKS)

## Notes

### Competing Interest Statement

The authors have declared no competing interest.

